# Phenotypic and Functional Characterization of Oncohistone Mutations in Breast Cancers

**DOI:** 10.1101/2025.04.29.651098

**Authors:** Andrea D. Edwards, Yangyang Dai, Siddharth Singh, Micah Thornton, Tulip Nandu, Ralf Kittler, Cristel V. Camacho, Dan Huang, W. Lee Kraus

## Abstract

Although mutations in genes encoding histones display a similar prevalence to that of some other somatic mutations in cancer, the underlying mechanisms by which histone mutations drive tumorigenesis have not been fully explored. Herein, we curated missense mutations occurring in core histone genes in breast cancers using data from MSK-IMPACT and cBioPortal to identify high frequency breast cancer-associated histone gene mutations. We characterized 17 high frequency oncohistone mutations in H2A H2B, and H3 that are enriched in breast cancer samples and occurred at glutamate (E), aspartate (D), serine (S), and arginine (R) residues. Many of these mutants co-occur with *PIK3CA* mutations in breast cancer samples. The oncohistone mutants were expressed in MCF-7 breast cancer cells or MCF-10A mammary epithelial cells and screened for effects on oncogenic phenotypes in a variety of cell- and tumor-based assays (i.e., proliferation, migration, invasion, competitive outgrowth, transformation). In addition, we examined the effects of selected mutants on DNA damage and gene expression. Our results indicate that the collection of oncohistone mutants that we screened have varying phenotypic and functional effects. Some can promote cancer-related phenotypes, with H2B-E76Q, H3-E97K, and H3-E105K eliciting strong oncogenic phenotypes and alterations in gene expression. All three of these mutants showed cooperativity with an activating mutation in PIK3CA (E545K), or a chemical activator of PI3Kα (UCL-TRO-1938), in assays of MCF-10A proliferation. H3-E105K also strongly promoted transformation of MCF-10A cells in an assay of growth on low attachment substrate independently of PIK3CA. Our results indicate that some high frequency oncohistone mutants can have oncogenic activity in breast cancers, and may act as potential cancer drivers. Collectively, these observations presented here can be used as a resource to connect biological and molecular outcomes to oncohistones in breast cancers.

**Significance:** In this study, we identified and characterized 17 high frequency oncohistone mutations in H2A H2B, and H3 that are enriched in breast cancer samples and promote cancer-related phenotypes when expressed in cells. Many of these oncohistone mutations co-occur in breast cancer samples with mutations in *PIK3CA,* the most frequently mutated gene in breast cancer, and may act as potential cancer drivers.

## Introduction

Breast cancer is a deadly disease that affects about 1:8 women during their lifetimes [1, 2]. The molecular etiology of breast cancer includes both genetic and epigenetic abnormalities [2, 3], including amino acid-altering mutations in genes encoding core histones [4, 5], that can have a significant impact on pathogenesis. Histones are an integral component of chromatin, a protein-DNA structure comprised of nucleosomes that is essential for maintaining genome integrity and functions, including epigenetic regulation, DNA repair, replication, and transcription [6–8]. The functions of histones within nucleosomes are regulated by post-translational modifications (PTMs), such as methylation, phosphorylation, and acetylation, which occur in the amino-terminal tails and histone fold domains [9]. Interestingly, somatic mutations in histones, which are present in 4% of diverse tumor types, occur at or near post-translational modification sites [4, 5, 10]. For example, the first reported somatic histone mutations occurring at H3.3-K27M in pediatric brain tumors [11, 12] and H3.3-K36M in chondroblastomas [13] block histone methylation at those sites and promote cancer phenotypes. Although histone mutations display a similar genomic prevalence to that of other somatic mutations in cancer, the underlying mechanisms by which other histone mutations drive tumorigenesis are not well characterized [5, 10].

ADP-ribosylation (ADPRylation) is a reversible PTM resulting in the covalent attachment of ADP-ribose (ADPR) units derived from β-NAD^+^ on a variety of amino acid residues in substrate proteins (e.g., Glu, Asp, Ser) [14, 15]. ADPRylation may result in the attachment of monoADPR (MAR) or polymers of ADPR (polyADPR or PAR) on substrate proteins [14, 15]. ADPRylation is catalyzed by PARP family of poly(ADP-ribosyl) polymerases (PARPs) and mono(ADP-ribosyl) transferases, consisting of 17 members that have distinct structural domains, activities, subcellular localizations, and functions [14, 15]. PARP1 and PARP2, the major nuclear PARP enzymes, catalyze poly(ADP-ribosyl)ation (PARylation) on a variety of nuclear substrates, including histones [16, 17]. Initial studies of ADPRylation and PARPs were focused on the role of PARP1 in DNA repair [18], but our understanding of the role of PARPs in different biological processes has grown considerably [15, 19]. A number of seminal studies on PARP-mediated ADPRylation over the past decade have shifted the focus to the regulation of gene expression [20–23]. In this regard, PARP1 and PARP2 bind to chromatin at key regulatory regions in the genome [23–29]. New findings have demonstrated diverse roles for these PARPs in chromatin regulation, transcription, and RNA biology [14, 21, 23, 30].

Histones are a prominent acceptors of ADPR in the cell, which can occur at glutamate (E), aspartate (D), serine (S), and arginine (R) residues [4, 31–33]. Recent advances in chemical biology and mass spectrometry have led to the identification and functional analysis of specific sites of ADPRylation on histones [16, 33–35]. This includes S residues during genotoxic stress [36], E and D residues in response to DNA damage [34], E and D residues under physiological conditions such as adipogenesis [33], and R residues in cell proliferation [37]. Recently, we found that up to 60% of the most frequently-occurring core histone mutations in over 180 cancer types are at E and D residues [4, 38]. From these potential oncohistone mutants, we characterized the biological, biochemical, and molecular mechanisms of H2B-D51N/A and H4-D68N/A mutations in breast and ovarian cancer cell lines. The mutant histones exhibited reduced levels of ADPRylation and promoted proliferation when expressed in cells [4]. Cells expressing the H2B-D51 mutants also exhibited increased p300-mediated acetylation of H2B at many lysine residues, altered chromatin accessibility at enhancers and promoters, and altered patterns of gene expression. In contrast, cells expressing the H4-D68 mutants showed no discernible changes in gene expression outcomes, but exhibited higher genomic instability, supporting a pro-mutagenic phenotype [4].

Herein, we have further explored the role of oncohistones in breast cancer biology. We have curated a set of 17 histone mutations that occur at potential sites of ADPRylation (E, D, S, and R residues) and are prevalent in breast cancers. We have screened for oncogenic phenotypes and explored potential molecular mechanisms of action of these mutant histones. Our study sheds new light on the role of oncohistones in breast cancers, which may suggest new therapeutic strategies.

## Results

### High frequency breast cancer-associated histone mutations are enriched at glutamate and arginine residues

We previously examined the types and frequencies of histone mutations across a spectrum of cancer types [4]. Here, we designed a study that would allow for identification of high frequency breast cancer-associated histone mutations, followed by phenotypic and functional analyses of selected histone mutants in breast cancer cells (Fig. 1A). We curated missense mutations in core histone genes in breast cancers using the MSK-IMPACT clinical sequencing cohort [39] and several other published studies in cBioPortal [40]. These analyses revealed 364 breast cancer-associated histone mutations. We limited our analyses to 217 patient tumor samples harboring 236 total histone mutations (accounting for a few patients that had 2 or more mutations) with a tumor mutation burden (TMB) less than 10 mutations per mega base to increase the likelihood of identifying driver mutations (Fig. 1B; Fig. S1). Our analyses demonstrated that nearly 40% of all histone mutations occurred at glutamate (E) and arginine (R) residues (Fig. 1C). Both of these amino acids are known to be major acceptor sites for ADPR, along with serine (S) and aspartate (D) [31, 32]. To narrow down the top mutations, we identified those that were recurrent (>1 count) in the 217 patient tumor samples, resulting in a curated set of 17 histone mutations, including those that occur at a E (10), R (4), D (2), and S (1) in histones H2A, H2B, and H3 (Fig. 1, D and E; Fig. S1). Many of these residues are located in the histone fold domains of the nucleosome core (Fig. 1F). Mutation at some of these sites have been shown in biochemical and biophysical assays to affect nucleosome sliding (e.g., H2A-E92 and -E121; H2B-E113) and destabilize nucleosomes (e.g., H2B-E71 and -E76; H3-E50 and -E97) [41]. Collectively, these results identify a high frequency set of breast cancer-associated histone mutations that we expected to alter outcomes in phenotypic and functional assays.

**Figure 1.**
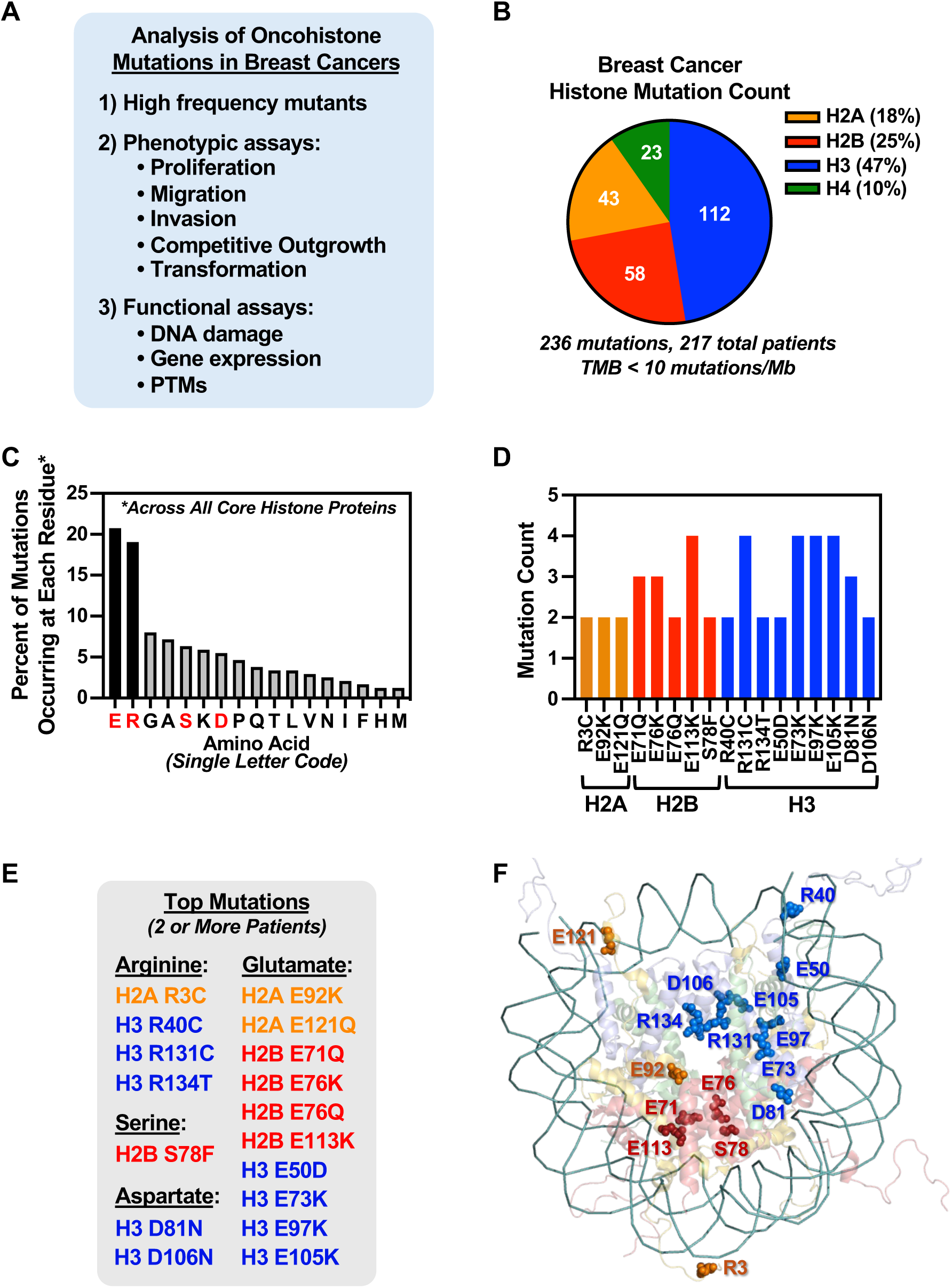
Histone mutations in breast cancers are enriched for ADPR acceptor residues. **(A)** Overall scheme for the identification and functional analysis of potential oncohistone mutants in breast cancers. **(B)** Pie chart showing the distribution of 236 histone mutations across H2A (*orange*), H2B (*red)*, H3 (*blue*) and H4 (*green*) occurring in a total of 217 breast cancer patient samples. Samples with a tumor mutation burden (TMB) of <10 mutations per mega base were selected for analysis. **(C)** Bar graph showing the frequency (in percentage) of mutation occurring at the specified amino acid residues. ADPR acceptor residues (E, R, S, and D) are highlighted in red. **(D)** Bar graph showing the top 17 histone mutations selected for further analysis based on mutation count (≥2) and amino acid residue of interest (E, R, S, and D). Bar colors indicate H2A (*orange*), H2B (*red)* and H3 (*blue*). **(E and F)** Location of histone mutations in the nucleosome structure. (E) List of the specific mutations highlighted in the nucleosome structure shown in panel (F).

### Evaluation and selection of breast cancer cell lines for phenotypic and functional assays of oncohistone mutants

In order to study the molecular, cellular, and biological effects of each oncohistone mutant, we evaluated and selected breast cancer cell lines for our analyses. We first examined the repertoire of existing histone gene mutations for a set of eight commonly used breast cancer cell lines. To do this, we performed targeted histone gene sequencing on cDNAs generated from the cell lines using a pool of primer pairs that amplifies all 61 core histone genes (Fig. S2). None of the 17 histone gene mutations of interest were found in any of the cell lines, although we did observe single nucleotide polymorphisms (SNPs) at some sites of interest that do not change the amino acids (Fig. S2A). We did find missense mutations at other residues, some of which are found in human cancers (Fig. S2B; H3-M72I and -M72V, H4-V3A). Excluding the top 5 mutations frequently found across all of the cell lines that we sequenced, MCF-7 cells harbored only two additional mutations (H2B-E3D and H2B-I40M) (Fig. S2B). By this analysis, MCF-7 cells are a useful system to reduce possible cooperativity from other pre-existing histone gene mutations.

With regard to cooperativity, histone gene mutations frequently co-occur with other somatic mutations at later stages of tumor progression [42]. *PIK3CA*, the gene encoding phosphatidylinositol-4,5-bisphosphate 3-kinase catalytic subunit alpha (PIK3CA), is the most frequently mutated gene in breast cancer (∼20-40% of breast cancers) [43–45]. Phosphatidylinositol 3′-kinases are lipid kinases that link oncogenes to multiple receptor-mediated signaling pathways (e.g., AKT) to drive cancer cell phenotypes [46, 47]. Activating mutations in *PIK3CA* enhance signaling and drive breast cancer phenotypes [45, 46, 48]. Moreover, activating mutations in *PIK3CA* may cooperate with other drivers, including oncohistones. For example, H2B-E76K enhances colony formation in the presence of PIK3CA H1047R [42]. We found that *PIK3CA* mutations occur in 47% of the breast cancer samples containing somatic histone gene mutations that we curated (Table 1A). Additionally, 12 out of 17 (70%) of the samples harboring the histone gene mutations selected for further analysis have co-occurring *PIK3CA* mutations (Table 1B).

**Table 1.** Frequency of co-occurring mutations with oncohistone mutations in breast cancers.

| Co-Occurring Mutation | No. Samples with the Mutation | Percent |
| --- | --- | --- |
| *PIK3CA* | 93 | 47 |
| *TP53* | 83 | 41 |
| *TTN* | 65 | 39 |
| *MUC16* | 39 | 25 |
| *KMT2C* | 34 | 17 |
| *RYR2* | 34 | 16 |
| *H3C2* | 32 | 16 |
| *SYNE1* | 25 | 15 |
| *SYNE2* | 22 | 14 |
| *NEB* | 31 | 11 |
| *SPEN* | 31 | 10 |

| Histone | Histone Mutation | *PIK3CA* Mutation |
| --- | --- | --- |
| H2A | H2A R3C | ☐ |
|  | H2A E92K |  |
|  | H2A E121Q | ☐ |
| H2B | H2B E71Q | ☐ |
|  | H2B E76K | ☐ |
|  | H2B E76Q |  |
|  | H2B E113K | ☐ |
|  | H2B S78F | ☐ |
| H3 | H3 R40C |  |
|  | H3 E50D | ☐ |
|  | H3 E73K | ☐ |
|  | H3 D81N |  |
|  | H3 E97K | ☐ |
|  | H3 E105K | ☐ |
|  | H3 D106N |  |
|  | H3 R131C | ☐ |
|  | H3 R134T | ☐ |

Presence or absence of a co-occurring *PIK3CA* mutation in tumor samples containing the top 17 histone mutations that were selected for analysis. A total of 12 out of 17 samples had co-occurring histone and *PIK3CA* mutations.

Given that MCF-7 cells have been reported to harbor a heterozygous *PIK3CA* E545K mutation [49], coupled with the lowest burden of pre-existing histone mutations in the cell lines that we screened, we chose MCF-7 cells for further evaluation of the selected histone mutants. Our goal was to examine a broad range of phenotypic outcomes for cells expressing the different histone mutants, including proliferation, migration, invasion, outgrowth, and transformation.

### Effect of oncohistone mutants on cell proliferation

We generated MCF-7 cells lines with stable and constitutive ectopic expression of individual FLAG-tagged histones, including wild-type H2A, H2B, or H3, or each of the 17 different H2A, H2B, or H3 mutants, as we described previously [4] (Fig. 1E). This system allows for ectopic expression at about 15% of the level of the cognate endogenous histone [4]. For the mutants, we used the naturally occurring amino acid alteration (Fig. 1E). We confirmed the roughly equal expression by blotting for FLAG (Fig. 2A). When then used these cell lines in proliferation assays to determine the effects of the histone mutants on proliferation in two-dimensional (2D) cell culture assays. MCF-7 cells ectopically expressing a wild-type or mutant histone were plated individually and cell proliferation was monitored over a period of 7 days. We observed that 3 of the mutants out of the 17 tested significantly enhanced cell proliferation compared to their respective wild-type controls: H2B-E76Q, H2B-E76K, and H3-R134T (Fig. 2B). Interestingly, expression of H2B-E113K significantly decreased proliferation of MCF-7 cells. These results indicate that some oncohistone mutants can enhance cell proliferation.

**Figure 2.**
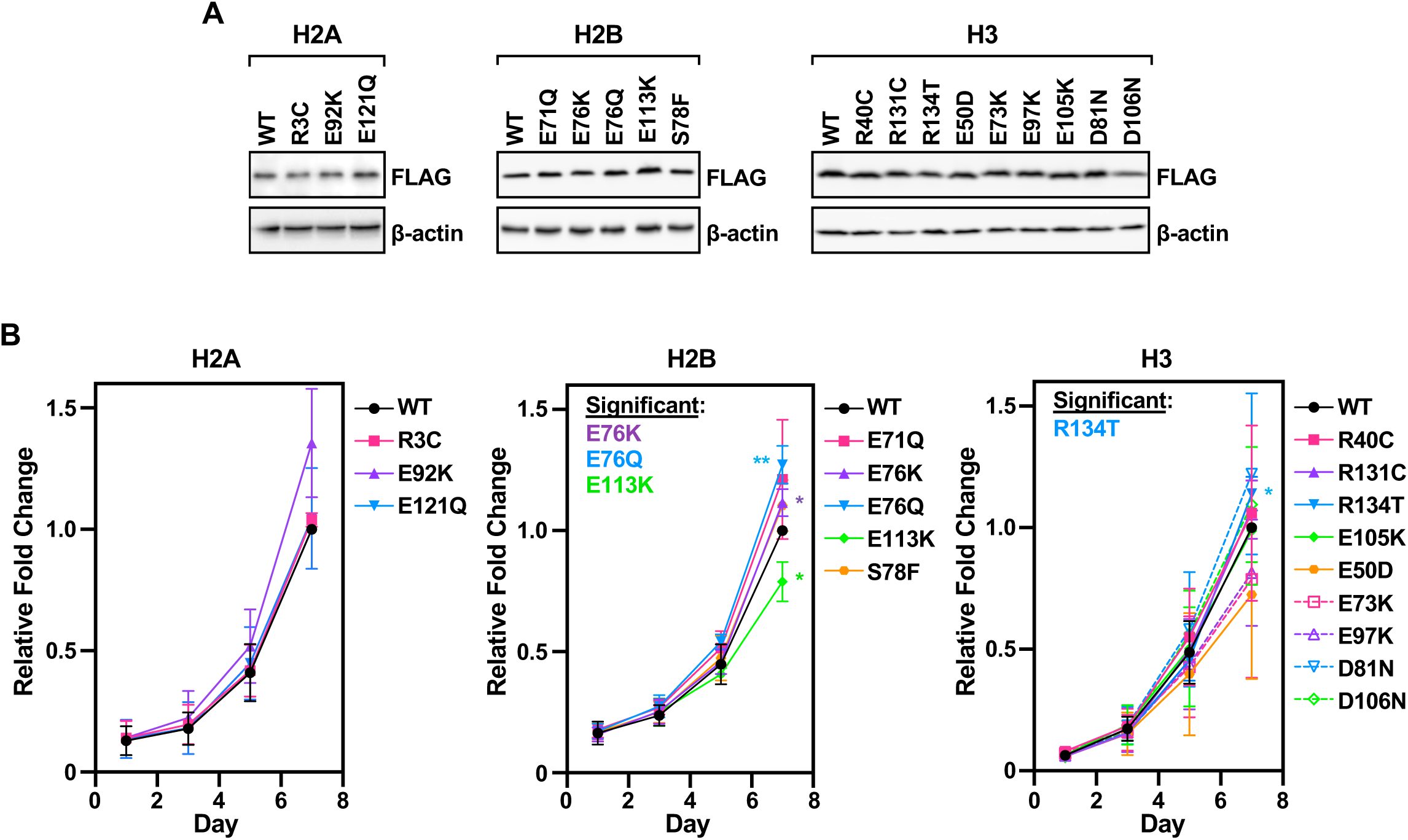
Proliferation of MCF-7 cells expressing histone mutants. **(A)** Ectopic expression of histone mutants in MCF-7 cells. Western blots showing the levels of ectopically expressed FLAG-tagged core histone proteins in extracts from MCF-7 cells. β-actin is shown as a loading control. **(B)** Line graphs showing the growth of MCF-7 cells ectopically expressing FLAG-tagged oncohistone mutants assayed by crystal violet staining. Each point represents the mean ± SEM; n = 3. Asterisks indicate the significance of the differences between wild-type and mutant at day 7; Unpaired t-test; * p < 0.05, ** p < 0.01. The four histone mutants that with significant effects are highlighted in the callout.

### Effect of oncohistone mutants on cell migration and invasion

Many phenotypes contribute to tumorigenicity and aggressiveness of tumors in vivo. Therefore, we sought to expand our screen beyond proliferation by examining cell migration and invasion. The MCF-7 cells ectopically expressing individual wild-type or mutant histones described above were plated individually in Boyden chambers to test for migration (without Matrigel) or invasion (with Matrigel). For cell migration, we found that 6 mutants significantly enhanced cell migration compared to their respective wild-type controls: H2A-E92K, H2A-E121Q, H2B-E76K, H2B-E76Q, H3-E73K, and H3-E97K (Fig. 3, A and B; Fig. S3).

**Figure 3.**
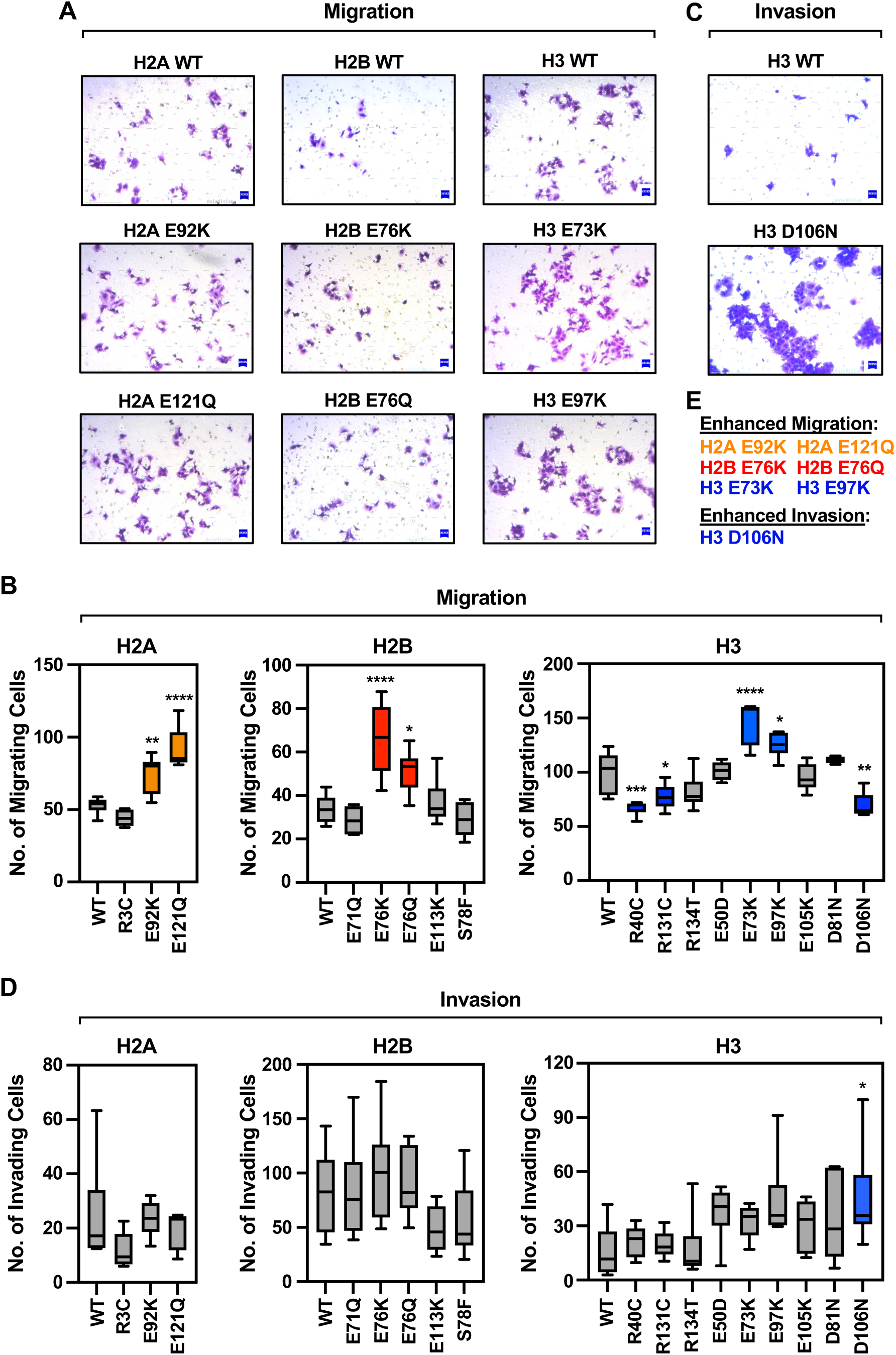
Migration and invasion of MCF-7 cells expressing histone mutants. Cell migration (A and B) and invasion (C and D) assays were performed in MCF-7 cells ectopically expressing FLAG-tagged histone mutants, as shown in Figure 2A. **(A and B)** Cell migration assays in MCF-7 cells ectopically expressing FLAG-tagged histone mutants, as indicated. (A) Representative images of migrating MCF-7 cells expressing individual histone H2A, H2B, and H3 mutants versus the corresponding wild-type histone. The cells were stained with crystal violet. Only statistically significant results are shown [shown as colored boxes in panel (B)]. (B) Box plots showing quantification of migrating cell numbers. H2A, n = 3; H2B, n = 4; H3, n = 4. Asterisks indicate significant differences between the mutant and wild-type; One-way ANOVA; * p < 0.05, ** p < 0.01, *** p < 0.0005, **** p < 0.0001. Colored boxes indicate mutants that had statistically significant effects versus wild-type. **(C and D)** Invasion assays in MCF-7 cells ectopically expressing FLAG-tagged histone mutants, as indicated. (C) Representative images of migrating MCF-7 cells expressing individual histone H3 mutants versus the corresponding wild-type histone. The cells were stained with crystal violet. Only statistically significant results are shown [shown as a colored box in panel (D)]. (D) Box plots showing quantification of invading cell numbers. H2A, n = 3; H2B, n = 4; H3, n = 3. Asterisks indicate significant differences between the mutant and wild-type; One-way ANOVA; * = p < 0.05. **(E)** List of oncohistone mutations showing significant effects on cell migration and invasion.

Interestingly, the H3 mutants R40C, R131C, and D106N significantly decreased cell migration compared to the wild-type H3 control. Overall, the histone mutants did not appear to have much impact on invasion (Fig. 3, C through E; Fig. S4), but the H3-D106N mutant significantly increased cell invasion compared to the wild-type H3 control (note that the results with the H3-E97K mutant approached significance) (Fig. 3, C and D). The observed effects with H3-D106N (inhibition of migration, enhancement of invasion) demonstrate that histone mutants can act differently on different phenotypic outcomes. The limited effects of the histone mutants on invasion in these assays might be attributable to the intrinsic nature of the mutation itself or due to the absence of the appropriate cellular microenvironment needed to reveal the invasion potential.

### Effect of oncohistone mutations on tumor growth in vivo

In order to identify oncohistone mutations with driver potential (versus passenger mutations), we performed competitive three dimensional (3D) outgrowth experiments using xenograft tumors formed in mice from pools of MCF-7 cells ectopically expressing individual FLAG-tagged wild-type or mutant histones (as shown in Fig. 2A). The cell lines expressing each of 17 histone mutants that are the focus of our experiments, or the corresponding wild-type histones, were pooled in equal numbers by individual histone (H2A, H2B, or H3) or all three together (All) and used to generate xenograft tumors in immunocompromised mice (Fig. 4A). At 40 days, we observed a significant growth of tumors formed from (1) the H2A mutant pool compared to wild-type H2A and (2) the all-histone pool compared to parental MCF-7 cells (Fig. 4B). At the end of the experiment, we performed targeted histone gene sequencing on cDNAs generated from the tumors using a pool of primer pairs that amplifies the ectopically expressed FLAG-tagged core histone genes. This allowed us to determine if any of histone genes were enriched in the population over time versus the starting pool. In each pool, we observed significant enrichment of a single histone mutant which conferred a growth advantage on the cells expressing it: E121Q for H2A, E76Q for H2B, and E105K for H3 and the all-histones pool (Fig. 4, C and D). These results indicate that H3-E105K is highly tumorigenic and confers a selective growth advantage in a pool of histone mutations in vivo.

**Figure 4.**
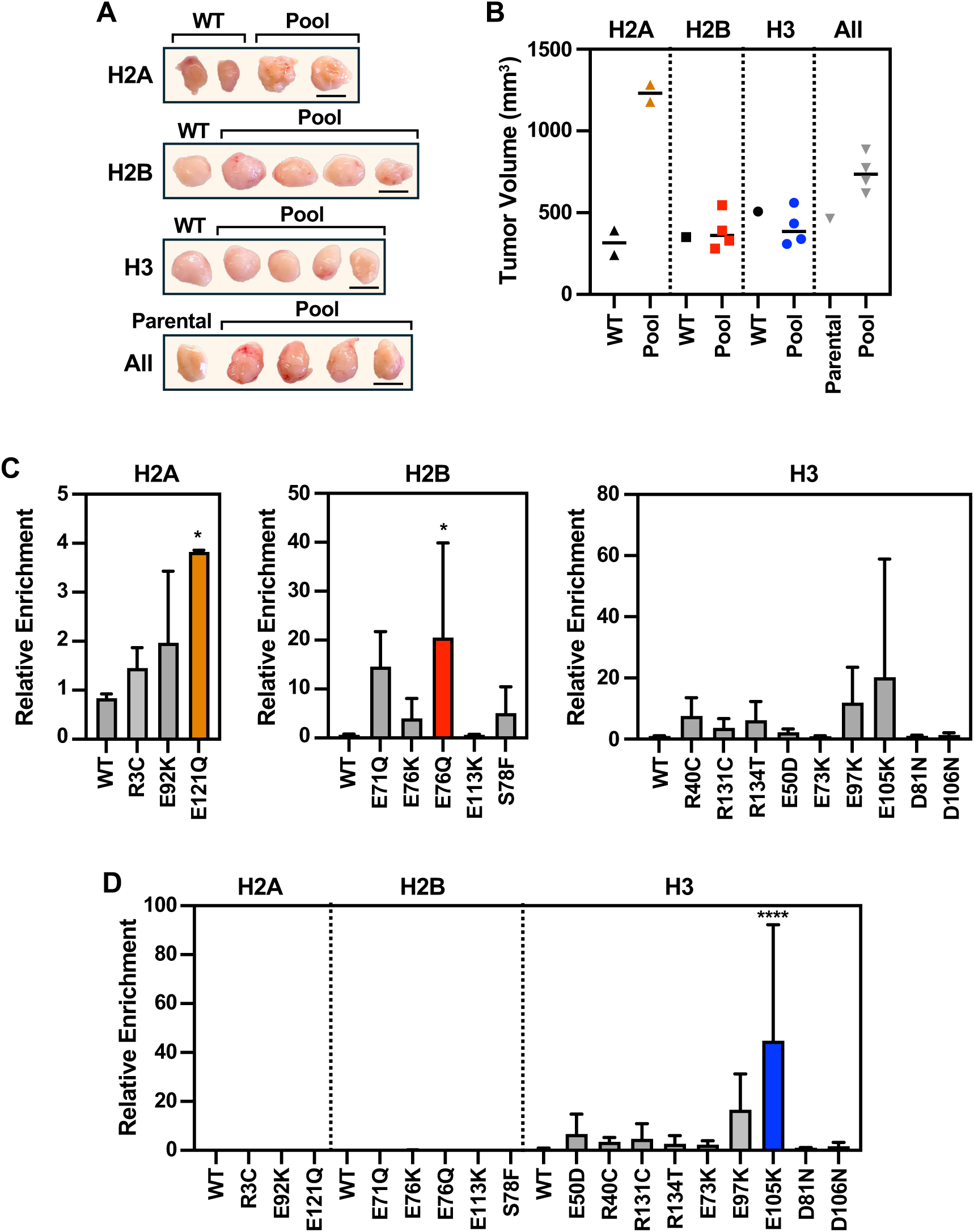
Competitive growth of MCF-7 cells expressing histone mutants in xenograft tumors. Xenograft assays in mice were performed using MCF-7 cells ectopically expressing FLAG-tagged histone mutants, as shown in Figure 2A. The tumors were formed by pooling the cells expressing the histone mutants in equal proportions. **(A)** Morphology of the xenograft tumors formed from pools of cells expressing mutant H2A, H2B, H3, a mix of all three, or parental MCF-7 cells collected at day 40 post-injection. Scale bars = 10 mm. **(B)**. Graph showing the final tumor volumes of the MCF-7 xenograft tumors from panel (A). **(C)** Bar graphs showing the relative enrichment of cells expressing each histone mutant in the xenograft tumors from (A) after sequencing using primers designed to amplify the ectopically expressed histones genes. Histone mutant genes were quantified and normalized to total reads per sample. Each bar represents the mean + SEM; H2A pool, n = 2; H2B pool, n = 4; H3 pool, n = 4. Asterisks indicate significance; One-way ANOVA; * p < 0.05. **(D)** Bar graph showing the relative enrichment of cells expressing each histone mutant in xenograft tumors containing all histone mutants from (A). Histone mutant genes were quantified and normalized to total reads per sample. Each bar represents the mean + SEM; Pool, n = 4. Asterisks indicate significant difference compared to wild-type; One-way ANOVA; **** p<0.0001.

### Effects of oncohistone mutations on DNA damage signaling and genomic stability

Our previous study demonstrated that breast cancer cells expressing H4-D68A/N mutants exhibited increased DNA damage under basal growth conditions as determined by the presence of γH2AX foci, suggesting higher genomic instability [4]. Therefore, we examined if the mutant histone identified herein could modulate DNA damage signaling and genomic stability. MCF-7 cells ectopically expressing wild-type or mutant histones were grown under basal conditions and stained with an antibody against phosphorylated H2AX (S139; γH2AX) to monitor DNA damage (treatment with H_2_O_2_ was used as a positive control). Most of the histones that elicited significant effects in this assay reduced the number of visible foci compared to their respective wild-type controls: H2B-E71Q, H2B-E76K, H2B-E113K, H2B-S78F, and H3-E73K (Fig. S5). This may indicate loss of proper DNA damage signaling. In contrast, H2A-R3C and H2A-E92K showed a modest, but significant increase in DNA damage foci, indicating higher genomic instability (Fig. S5). Collectively, these results indicate that effects on DNA damage may not be the predominant mechanism driving phenotypic outcomes with oncohistone mutants.

### Cooperative effects of selected H2B and H3 mutants with an activating PIK3CA mutant on proliferation and transformation

Following up on our initial results, we determined if selected oncohistone mutants that exhibited oncogenic phenotypes in the assays described above can drive transformation of the non-transformed MCF-10A human mammary epithelial cells in cooperation with the activating PIK3CA mutation E545K. For these assays, we focused on H2B-E76Q, H3-E97K, and H3-E105K, which showed strong phenotypes in one or more of the assays described above. Mutations at H2B-E76 have previously been shown to elicit cancer-related phenotypes, alter gene expression, and drive transformation [41, 42, 50]. H2B-E76Q and H3-E97K are located in the histone fold domain and have been shown to decrease nucleosomes stability [41]. H3-E105K has not been studied in detail. Mutant PIK3CA is co-expressed with all three histone mutants in breast cancers (Table **1**).

We ectopically expressed the three histone mutants (i.e., H2B-E76Q, H3-E97K, and H3-E105K), or their cognate wild-type histones (i.e., H2B and H3), individually in an MCF-10A cell line expressing the PIK3CA-E545K mutant [51] (Fig. 5A). All three oncohistone mutants promoted cell proliferation compared to their cognate wild-type histones (Fig. 5, B and C). Additional assays of growth on low attachment culture dishes, which provides an indication of transformation [52], showed that H2B-E76Q and H3-E105K, but not H3-E97K, significantly enhanced cell sphere formation compared to their cognate wild-type histones (Fig. 5D). The results with H2B-E76Q in these assays are similar to those reported previously for H2B-E76K [50]. Together these results indicate that (1) these three oncohistone mutants cooperate with PIK3CA-E545K and (2) H2B-E76Q and H3-E105K can cooperate with PIK3CA-E545K to drive transformation of MCF-10A cells.

**Figure 5.**
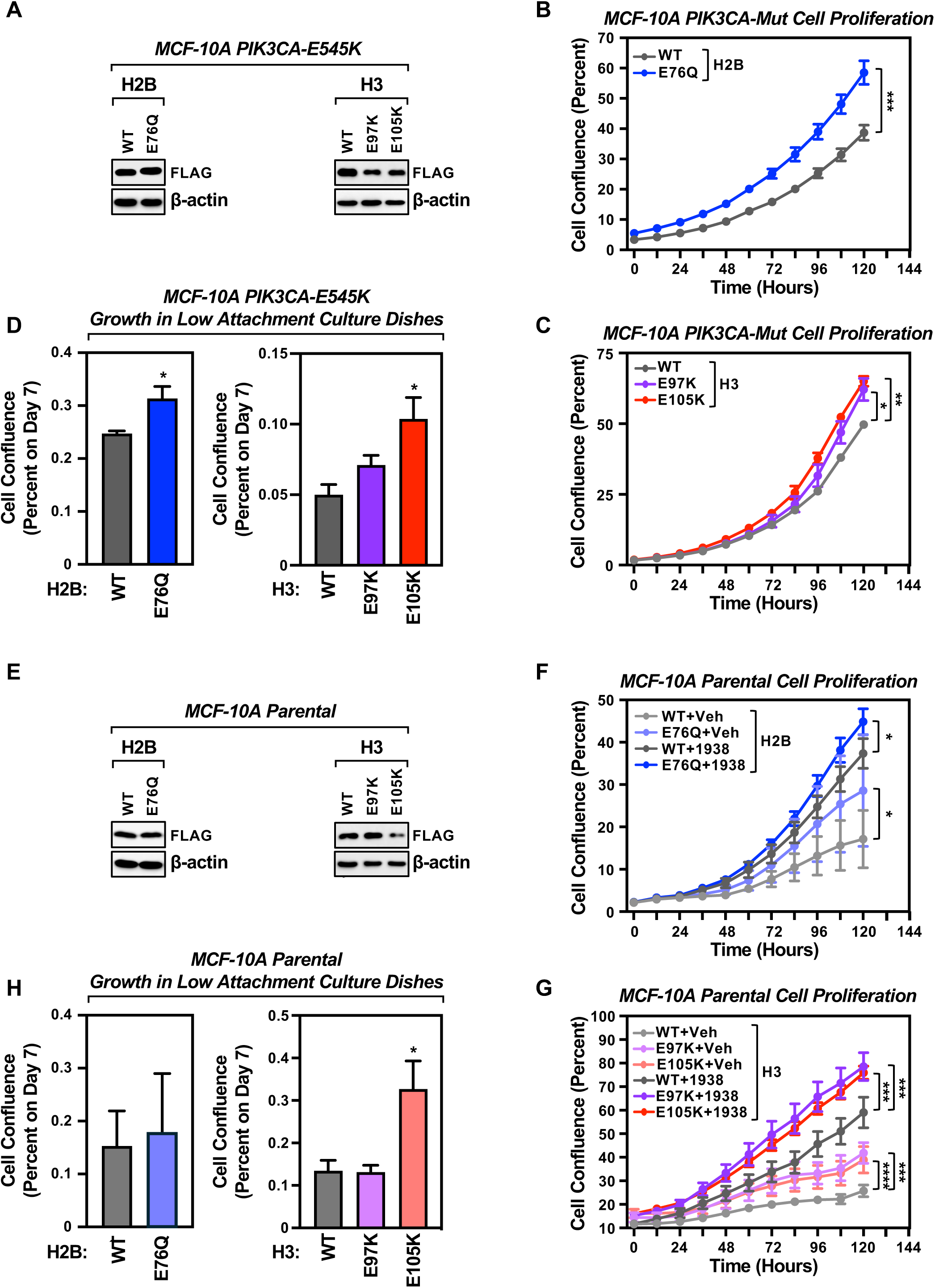
Cooperative effects with an activating PIK3CA mutant, as well as individual effects, on proliferation and transformation for three selected H2B and H3 mutants. Proliferation and transformation assays for selected oncohistone mutants (H2B-E76Q, H3-E97K, and H3-E105K) that exhibited oncogenic phenotypes in cell-based assays. We ectopically expressed the three histone mutants, or their cognate wild-type histones, individually in MCF-10A cells expressing the PIK3CA-E545K mutant [51] (A through D) or in parental MCF-10A cells (E through H). **(A)** Ectopic expression of three selected histone mutants in MCF-10A cells expressing the PIK3CA-E545K mutant. Western blots showing the levels of ectopically expressed FLAG-tagged core histone proteins in extracts from MCF-10A PIK3CA-E545K mutant cells. β-actin is shown as a loading control. **(B and C)** Line graphs showing the proliferation assay for MCF-10A cells co-expressing the PIK3CA-E545K mutant and individual FLAG-tagged histones mutants using an IncuCyte imaging system. (B) H2B-E76Q, (C) H3-E97K and H3-E105K. Each point represents the mean ± SEM; n = 3. Asterisks indicate the significance of the differences between wild-type and mutant at day 5; Unpaired t-test; * p < 0.05, ** p < 0.01, *** p < 0.001. **(D)** Bar graphs showing the growth of MCF-10A cells co-expressing the PIK3CA-E545K mutant and individual FLAG-tagged wild-type or mutant histones on low attachment culture dishes. *(Left)* H2B-E76Q, *(Right)* H3-E97K and H3-E105K. Each point represents the mean ± SEM; n = 3. Asterisks indicate the significance of the differences between wild-type and mutant at day 7; Unpaired t-test; * p < 0.05. **(E)** Ectopic expression of three selected histone mutants in parental MCF-10A. Western blots showing the levels of ectopically expressed FLAG-tagged core histone proteins in extracts from MCF-10A cells. β-actin is shown as a loading control. **(F and G)** Line graphs showing the proliferation assay for MCF-10A cells expressing individual FLAG-tagged wild-type or mutant histones assayed using an IncuCyte imaging system. (F) H2B-E76Q, (G) H3-E97K and H3-E105K. Cells were treated with 5 μM of the PI3K activator UCL-TRO-1938 or DMSO vehicle. Each point represents the mean ± SEM; n = 3. Asterisks indicate the significance of the differences between wild-type and mutant at day 5; Unpaired t-test; * p < 0.05, ** p < 0.01, *** p < 0.001, **** p < 0.0001. **(H)** Bar graphs showing the growth of MCF-10A cells expressing individual FLAG-tagged wild-type or mutant histones on low attachment culture dishes. *(Left)* H2B-E76Q, *(Right)* H3-E97K and H3-E105K. Each point represents the mean ± SEM; n = 3. Asterisks indicate the significance of the differences between wild-type and mutant at day 7; Unpaired t-test; * p < 0.05.

**Figure 6.**
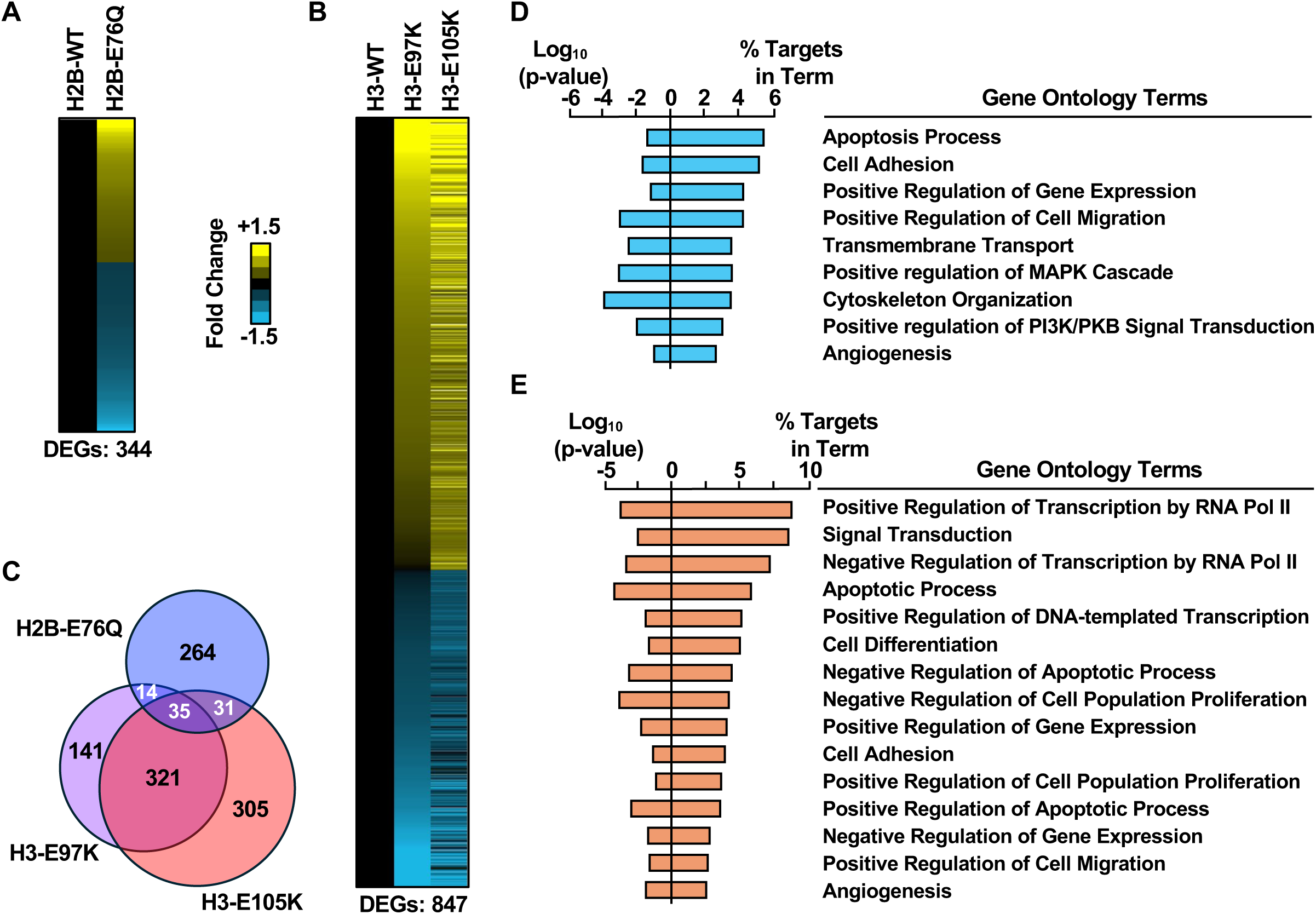
Gene regulatory effects of selected H2B and H3 mutants. Gene expression and gene ontology (GO) analyses for selected oncohistone mutants (H2B-E76Q, H3-E97K, and H3-E105K) that exhibited oncogenic phenotypes in cell-based assays. We ectopically expressed the three histone mutants, or their cognate wild-type histones, individually in MCF-7 and performed genomic and bioinformatic analyses. **(A and B)** Heatmaps representing the fold change of differentially expressed genes (DEGs) from RNA-seq performed in MCF-7 cells ectopically expressing (A) FLAG-tagged wild-type H2B or H2B-E76Q or (B) FLAG-tagged wild-type H3, H3-E97K, or H3-E105K. Results are for mutant versus wild-type. **(C)** Venn diagrams showing the overlap of affected genes for each of the three histone mutants from (A) and (B). **(D and E)** Gene ontology terms enriched for differentially expressed genes in MCF-7 cells ectopically expressing (D) FLAG-tagged H2B-E76Q or (E) FLAG-tagged H3-E97K or H3-E105K. Results are for each mutant versus wild-type.

### Individual effects of selected H2B and H3 mutants on proliferation and transformation

To determine if the three selected oncohistone mutants could drive proliferation and transformation independently of mutations in PIK3CA, we assayed their effects in parental MCF-10A cells in which we ectopically expressed the histone mutants (i.e., H2B-E76Q, H3-E97K, and H3-E105K), or their cognate wild-type histones (i.e., H2B and H3) (Fig. 5E). As before, H2B-E76Q, H3-E97K, and H3-E105K promoted MCF-10A cell proliferation compared to the cognate wild-type histones (Fig. 5, F and G). Although treatment with the PI3K activator UCL-TRO-1938 [53] dramatically increased cell proliferation compared to the vehicle treatment with the wild-type and mutant histones, it did not alter the relative effects of the histone mutants compared to wild-type histones (Fig. 5, F and G). Additional assays of growth on low attachment culture dishes revealed that MCF-10A cells ectopically expressing H3-E105K, but not H2B-E76K or H2B-E76Q, significantly increased cell sphere formation (Fig. 5H). These results suggest that H3-E105K is able to drive cell transformation from a non-oncogenic to oncogenic phenotype.

### Gene regulatory effects of selected H2B and H3 mutants

Oncohistone mutations have been linked to regulation of gene expression [4, 41, 42, 50]. To determine whether the three selected H2B and H3 mutants could impact gene expression in MCF-7 cells, we performed RNA-sequencing in cells ectopically expressing individual mutants or the cognate wild-type histones. All three of the mutant histones altered gene expressed compared to the cognate wild-type histones (Fig. 7, A and B). The genes affected by the H3 mutants were more similar to each other than the those affected by the H2B mutant (Fig. 7C). Gene ontology (GO) analyses indicated that the genes significantly altered by the H2B-E76Q mutant were enriched in GO terms related to the regulation of gene expression, cell migration, cell adhesion, and apoptotic processes, as well as MAPK and PI3K/PKB signaling pathways (Fig. 7D), and that the genes significantly altered by the H3-E97K and H3-E105K mutants were enriched in GO terms related to the regulation of transcription, gene expression, cell proliferation, and cell migration (Fig. 7E). In general, these GO terms correspond to the cancer-related phenotypes observed with these oncohistone mutants.

## Discussion

In this study, we curated and explored the phenotypic and functional outcomes of high frequency oncohistone mutations in breast cancers. Our analyses included a sequence-based characterization of histone gene mutations in eight commonly used breast cancer cell lines, which should be a useful resource for future studies on histone mutations in breast cancers. We selected 17 recurrent histone mutants and characterized them for their potential oncogenic effects in breast cancer cells. Through a series of cell-based and xenograft experiments, we observed various oncogenic gain-of-function effects elicited by several selected oncohistone mutants (Fig. S6). For instance, H3-E105K conferred high tumorigenic potential, H2B-E76Q increased proliferation, H2A-E121Q, H2B-E76K and H3-E97K enhanced migration, and H3-D106N enhanced invasion. Of all the oncohistone mutants that we tested, H3-E105K had the greatest potential to drive transformation, with H2B-E76Q and H3-E97K showing some transformation potential. Surprisingly, a few of the oncohistone mutants (for example, H2B-E113K) elicited effects which could be characterized as tumor suppressive (Fig. S6). Three mutants that we selected for more detailed analyses (i.e., H2B-E76Q, H3-E97K, and H3-E105K) promoted alterations in gene expression when expressed in breast cancer cells. Some of the amino acids mutated in the oncohistones that we studies are putative sites for PTMs, such as ADPRylation or methylation. The modification of these sites and the functions of these PTMs will be assessed in future studies.

### Frequent occurrence of oncohistones mutations in the globular domains of histones

Among the selected oncohistones in this study, only three (out of seventeen) of the histone mutations (H2A-R3C, H2A-E121Q, and H3-R40C) were in the amino or carboxyl tail regions of the histones, while the others occurred in the residues within the globular, histone fold domains. Similar observations were made in previous studies, where the globular domains of the core histone proteins were revealed to be the major site of histone mutations, rather than the more accessible tail regions [38, 41]. Though several of the sites of histone core domain mutations are not known to be sites for PTMs, they can promote oncogenesis by altering nucleosome structure and stability, nucleosomes remodeling, or the modification of nearby residues [4, 38, 41, 50]. For instance, H2B-E76K has been previously shown to destabilize nucleosomes, with reduced interactions between H2A-H2B dimers and H3-H4 dimers [50, 54].

Recently, nucleosomes containing H3-E97K were shown to cause structural alterations near the mutant residue leading to reduced binding of linker histone H1, defective activation of PRC2, and enhanced transcription by RNA polymerase II [55]. Moreover, H2B-D51 was shown to dramatically alter chromatin accessibility at enhancers and promoters in cells, as well as promote p300-mediated acetylation of H2B at many lysine residues in H2B [4]. Hence, compelling evidence are present that indicates that globular domain mutations of histones can be detrimental for chromosomal architecture, with the potential to induce cancer-specific transcriptional changes.

### Oncohistone mutations promote different cellular phenotypes in breast cancer cells

In our curated set of 17 oncohistone mutants, some of mutants stood out as oncogenic in multiple assays (Fig. S6). These include H2A-E92K and H2A-E121Q; H2B-E76K and H2B-E76Q; and H3-E73K, H3-E97K, and H3-E105K, with H3-E105K having the greatest driver potential. In contrast, H3-R40C and H3-E113K showed tumor suppressive effects. We observed striking variations among the oncohistones tested in their ability to promote tumor cell phenotypes, such as proliferation, migration, invasion, tumorigenicity, and DNA damage induction. These differential biological effects conferred by different oncohistones in specific contexts, could be the eventual outcome of several mechanisms such as (1) structural perturbations upon mutation leading to altered histone octamer formation, changes in chromatin accessibility, and downstream gene expression changes that promote cancer [4, 54], (2) epigenomic alterations in the genome through *cis* or *trans* loss or gain of histone PTMs due to alterations in the function of histone modifying enzymes [4, 56], or (3) cooperativity of oncohistones with secondary mutations to promote oncogenesis such as in the case of H2B-E76K, which in synergizes with activating mutations in *PIK3CA* to promotes colony formation in MCF-10A cells [50], similar to what we observed with the H2B-E76Q mutant .

### Potential impact of charge alterations in oncohistone mutants on nucleosome structure

Histones are rich in positively charged (basic) lysine (K) and arginine (R) residues that are mostly concentrated in the tail region and some of these residues are critical for the interaction of histones with DNA, provide stability to the nucleosome by neutralizing the negative charge of the nucleosomal DNA [57]. Though negatively charged aspartate (D) and glutamate (E) residues in histones do not directly bind to DNA, they are potentially involved in fine tuning the interaction with DNA through electrostatic repulsion with the negatively charged phosphate backbone of DNA, and hence can influence chromatin structure and gene expression. In our curated set of 17 oncohistones, almost all have residue changes from glutamic acid (negatively charged/acidic) to lysine (positively charged/basic) or glutamine (uncharged) (Fig. 1, E and F), which could potentially disturb required charge repulsion and chromatin compaction. These mutations might also alter the interactions of nucleosomes with other biomolecules [58]. The majority of the histone mutations (10 out of 17) transformed the wild-type residues (positively/negatively charged) to an uncharged amino acid residue (Fig. 1, E and F). This again, can interfere with the charge balance needed for proper histone-DNA interaction and/or proper histone-histone interactions and octamer assembly [58].

### Potential impact of site alterations in oncohistone mutants on histone modifications

Site alterations in oncohistone mutants have the potential to destroy or create sites of posttranslational modification. For example, mutations in H3 in pediatric cancers (e.g., H3K27M in glioma [11, 12] and H3K36M in chondroblastoma [59]) disrupt sites of histone methylation and inhibit the downstream epigenomic effects [60–66]. We showed that H2B-D51 mutants alter chromatin-dependent gene regulation by disrupting a site of PARP1-mediated ADP-ribosylation (ADPRylation) and allowing unrestrained p300-mediated H2B acetylation [4]. Also, in a non-cancer system (i.e., preadipocytes), we demonstrated that mutation of H2B-E35, which disrupts a site of PARP1-mediated ADP-ribosylation, allows unrestrained phosphorylation of H2B-S36 by AMP-activated protein kinase (AMPK) [33]. These results suggest a functional interplay among oncohistone mutants, PTMs, and histone-modifying enzymes.

Collectively, our study has identified a rich landscape or oncohistone mutants in breast cancers. As we demonstrated in phenotypic and functional assays with selected mutants, oncohistones are capable of altering molecular processes that drive key cancer cell phenotypes.

## Methods

### Antibodies and specialized reagents

The other antibodies used were as follows: FLAG (Sigma-Aldrich, F3165; RRID:AB_259529), histone H2A (Abcam, ab18255; RRID:AB_470265), histone H2B (Abcam, ab1790; RRID:AB_302612), histone H3 (Abcam, ab1791; RRID:AB_302613), histone H4 (Abcam, ab10158; AB_296888), phospho-histone H2A.X (Ser139) antibody (Millipore; RRID:AB_309864), PAR binding reagent (Millipore, MABE1031; RRID:AB_2665467), pan-ADP-ribose binding reagent (Millipore, MABE1016; RRID:AB_2665466), β-actin (Cell Signaling, 3700; RRID:AB_2242334), goat anti-rabbit HRP-conjugated IgG (ThermoFisher, 31460), and goat anti-mouse HRP-conjugated IgG (ThermoFisher, 31430).

Secondary antibodies included goat anti-rabbit HRP-conjugated IgG (ThermoFisher, 31460; RRID:AB_228341), goat anti-mouse HRP-conjugated IgG (ThermoFisher, 31430; RRID:AB_10960845), rabbit IgG (ThermoFisher, 10500C; RRID:AB_2532981), and Alexa fluor 488 goat anti-mouse IgG (ThermoFisher; RRID:AB_2534069). For experiments with MCF-10A cells, we used PI3K activator UCL-TRO-1938 (MedChem Express, HY154848).

### Cell culture

MCF-7 cells were kindly provided by Dr. Benita Katzenellenbogen (University of Illinois at Urbana-Champaign, IL) and were maintained in minimal essential medium (MEM; Sigma M1018) supplemented with 5% calf serum (Sigma, C8056), 100 units/mL penicillin-streptomycin (Gibco, 15140122), and 25 μg/mL gentamicin (Gibco, 1571004). MCF-10A PIK3CA-E545K mutant cells [51] were kindly provided by Dr. Ben Ho Park (Vanderbilt University Medical Center, TN) and were maintained in DMEM/F12 (Invitrogen, 11330032) supplemented with 5% horse serum (Sigma, H1270), 10 μg/mL insulin (Sigma, I5500), 0.5 μg/mL hydrocortisone (Sigma, H4001), and 0.1 μg/mL cholera toxin (Sigma, C8052). All other cell lines were obtained from the American Type Culture Collection (ATCC): MCF-10A (RRID:CVCL_0598), ZR-75-1 (RRID:CVCL_0588), SK-BR-3 (RRID:CVCL_0033), HCC70 (RRID:CVCL_1270), MDA-MB-231 (RRID:CVCL_0062), HCC1937 (RRID:CVCL_0290), MDA-MB-468 (RRID:CVCL_0419), HEK-293T (RRID:CVCL_0063), and maintained as recommended. All cell lines were authenticated for cell type identity using the GenePrint 24 system (Promega, B1870), and confirmed as Mycoplasma-free every six months using the Universal Mycoplasma Detection Kit (ATCC, 30-1012K). Fresh cell stocks were regularly replenished from original stocks every few months (no more than 15 passages).

### Sequencing cell lines for endogenous histone mutations

Breast and ovarian cancer cell lines were cultured as described previously under basal conditions. Genomic DNA was isolated from each cell line using a genomic DNA isolation kit (BS88504, Bio Basic). Primer pairs were designed to capture all 61 of the histone genes from genomic DNA by multiplex amplification (the sequences of the primer pairs are available on request). Paired-end sequencing (300 nucleotides for each end) was performed using MiSeq. We obtained ∼22 million total reads per run or 2 million reads per sample. The Genomic Analysis Toolkit (GATK) software [67] was used to identify single nucleotide polymorphisms (SNPs) and missense mutations in reference to the human genome. Integrated Genomics Viewer (IGV) software [68]was used to sort through the mutations and identify locations on the chromosomes.

### Generation of lentiviral expression vectors for wild-type and site-mutant histones

Double-stranded cDNAs encoding carboxyl-terminal FLAG epitope-tagged human wild-type and site-mutant histones H3, H2B, H2A, and H4, were synthesized as gene blocks (Integrated DNA Technologies), and then cloned individually into pCDH-EF1α-MCS-IRES-Puro (System Biosciences) lentiviral expression vector using Gibson assembly (NEB, E2621). Seventeen individual arginine (Arg R), glutamate (Glu, E), aspartate (Asp, D) and serine (Ser, S) amino acids were changed to the naturally occurring mutant residue commonly found in human cancer databases.

### Generation of stable cell lines for ectopic expression of wild-type and site-mutant histones

Lentiviruses were generated by transfecting the pCDH vectors described above into 293T cells, together with (i) an expression vector for the VSV-G envelope protein (pCMV-VSV-G, Addgene 8454; RRID:Addgene_8454), (ii) an expression vector for GAG-Pol-Rev (psPAX2, Addgene 12260;RRID: Addgene_12260), and (iii) a vector to aid with translation initiation (pAdVAntage, Promega E1711) using Lipofectamine 3000 reagent (Invitrogen, L3000015) according to the manufacturer’s instructions. The resulting viruses were collected in the culture medium, concentrated using Lenti-X Concentrator (Clontech, 631231), and used to individually infect MCF-7, MCF-10A, or MCF-10A PIK3CA-E545K cells. Cells were selected with 2 μg/mL puromycin (Sigma-Aldrich, P9620), expanded, and frozen in aliquots for future use. Ectopic expression of the cognate proteins was confirmed by Western blotting.

### Preparation of whole cell lysates

Cells were collected, washed with ice-cold PBS, and resuspended in MNase Whole Cell Lysis Buffer (50 mM Tris-HCl pH 7.9, 2 mM CaCl_2_, 0.2% Triton X-100, 100 U/mL micrococcal nuclease (Worthington, LS004797), 1x complete protease inhibitor cocktail (Roche, 1169748001), 250 nM ADP-HPD (Sigma-Aldrich, A0627, a PARG inhibitor), 10 μM PJ34 (Enzo Life Sciences, ALX-270-289, a PARP inhibitor) and incubated for 15 min at 37°C with gentle mixing to lyse the cells and extract the proteins. After brief sonication, the lysates were clarified by centrifugation in a microcentrifuge for 5 min at 4°C full speed and cell extracts collected. Protein concentrations were measured using the Bio-Rad Protein Assay Dye Reagent and volumes of lysates containing equal total amounts of protein were mixed with 4x SDS loading dye with Bromophenol blue. The lysates were then incubated for 5 min at 100°C.

### Western blotting

Equal amounts of whole cell lysates were prepared and run on 15% polyacrylamide-SDS gels. The gels were then transferred to nitrocellulose membranes and blocked with 5% nonfat milk in Tris-Buffered Saline with 0.1% Tween 20 (TBST) for 1 hr at room temperature. Primary antibodies were diluted in TBST and incubated at 4°C overnight. The membranes were washed multiple times in TBST and then incubated with horseradish peroxidase (HRP)-conjugated secondary antibody diluted in 5% nonfat milk in TBST for 1 hr at room temperature. Signals were captured using a luminol-based enhanced chemiluminescence HRP substrate (SuperSignal™ West Pico, Thermo Scientific) or an ultra-sensitive enhanced chemiluminescence HRP substrate (SuperSignal™ West Femto, Thermo Scientific) and a ChemiDoc imaging system (Bio-Rad).

### Cell proliferation assays

Cell proliferation was assessed in MCF-7 cells using a crystal violet staining assay. The cells were plated at a density of 10^4^ cells per well in 12-well plates. The cells were collected every 2 days. After collections, the cells were washed with PBS, fixed for 10 min with 4% paraformaldehyde at room temperature, and stored in 4°C until all time points had been collected. The fixed cells were stained with 0.1% crystal violet in 20% methanol solution containing 200 mM phosphoric acid. After washing to remove unincorporated stain, the crystal violet was extracted using 10% glacial acetic acid and the absorbance was read at 595 nm. Cell proliferation was also assessed in parental and PIK3CA-E545K mutant MCF-10A cells. The cells were seeded at a density of 7.5 × 10^3^ cells/mL in 12-well plates. Parental MCF-10A cells were treated with 5 μM of the PI3K activator UCL-TRO-1938 (MCE, HY154848) or DMSO vehicle. Cell confluence was assessed every 12 hour using an IncuCyte imaging system (Sartorius). All proliferation assays were performed a minimum of three times using independent platings of cells to ensure reproducibility.

### Cell migration assays

Boyden chamber assays were used to determine the effects of histone mutations on cell migration. Cells were trypsinized, resuspended in serum-free media, and collected by centrifugation at 300 g for 3 min at room temperature. The cell pellets were then resuspended in serum-free media and 500 μL of media containing 100,000 MCF-7 cells were plated into the top chamber (Corning 353097). The chambers were then incubated in serum-containing media with calf-serum. The plates were incubated for 72 hours and then the inside of the chamber was gently wiped to remove the cells in the top chamber. The chamber was fixed with 4% PFA for 20 min then incubated in 0.5% crystal violet in 20% methanol for 20 min with gentle shaking and then washed with water. The chambers were air dried, and the membranes were photographed. Using the microscope, representative photos were taken of each chamber. The number of migrated cells was counted manually in each photograph. Three independent experiments were performed with independent biologic replicates. Ordinary one-way ANOVA tests were used to evaluate for significant differences between different treatments.

### Cell invasion assays

Cell invasion assays were performed similar to cell migration assay described above using migration chambers (Corning 354480). After 72 hours, the cells in the top chamber were removed. The chamber was fixed with 4% PFA for 20 min then incubated in 0.5% crystal violet in 20% methanol for 20 min with gentle shaking and then washed with water. The chambers were air dried, and the membranes were photographed. Using the microscope, representative photos were taken of each chamber. The number of invading cells was counted manually in each photograph. Three independent experiments were performed with independent biologic replicates. Ordinary one-way ANOVA tests were used to evaluate for significant differences between different treatments.

### Cell growth in low attachment culture dishes

To assay transformation potential, we used an assay to monitor cell growth under low attachment conditions [52]. Parental and PIK3CA-E545K mutant MCF-10A cells were seeded at a density of 1.2×10^4^ cells/mL in low attachment 3D cell floater dishes (SPL, 26100), and maintained for 7 days to allow the sphere formation. Cell sphere confluence was then assayed using the IncuCyte imaging system (Sartorius).

### Competitive cell growth in xenografts

#### Formation of xenograft tumors in mice

All animal experiments were performed in compliance with the Institutional Animal Care and Use Committee (IACUC) at the UT Southwestern Medical Center. Female NOD *scid* gamma (NSG) mice at 6–8 weeks of age were used. We used female mice because mammary cancers occur primarily in females. In addition, the human cancer cell lines that we used for xenografts are from females. Mice were supplemented with 1.7 mg 17β-estradiol in 60-day release pellets (Innovative Research, SE-121) implanted subcutaneously at the base of the neck under anesthesia. To establish breast cancer xenografts, 5 x 10^6^ MCF-7 cells engineered for ectopic constitutive expression were injected subcutaneously into the flank of the mice in a 1:1 ratio PBS and Matrigel (Fisher). Individual histone mutants were mixed in equal proportions. Pools included: (i) H2A pool; (ii) H2B pool; (iii) H3 pool; and (iv) all histones pool. Mouse weight was monitored once per week and tumor growth measured over time using electronic calipers approximately every 3-4 days. Tumor volumes were calculated using a modified ellipsoid formula: Tumor volume = ½ (*length* × *width*^2^). Animals were euthanized at 40 days post-injection. Fresh tumor samples were flash frozen in liquid nitrogen or fixed for 24 hours in 10% formalin.

#### Preparation of whole-cell lysates from xenograft tissues

Xenograft tissue was incubated in whole cell extract buffer (50 mM Tris pH 7.5, 0.5 NaCl, 0.5 M EDTA, 1% NP-40, and 10% glycerol) for 15 min on ice followed by maximum speed centrifugation for 30 min. The tumor tissues were homogenized to make extracts used for Western blotting. Protein concentrations were measured using the Bio-Rad Protein Assay Dye Reagent and volumes of lysates containing equal total amounts of protein were mixed with 4x SDS loading dye with Bromophenol blue. The lysates were then incubated for 5 min at 100°C.

#### Determination of cell outgrowth in xenograft tumors

Frozen tumor tissue was thawed on ice. Genomic DNA was isolated from ectopically expressing histone mutant tumors using a genomic DNA isolation kit (BioBasic BS88504). We used universal primers to capture and PCR-amplify ectopically expressed histone genes (the sequences of the primer pairs are available on request). For specificity to the ectopic histone genes, the reverse primer targeted the C-terminal FLAG sequence. Libraries were prepared from 10 ng DNA following the Illumina TruSeq protocol. The quality of the libraries was assessed using a D1000 ScreenTape on a 2200 TapeStation (Agilent) and quantified using a Qubit dsDNA HS Assay Kit (Thermo Fisher). Libraries with unique adaptor barcodes were multiplexed and sequenced on an Illumina MiSeq (paired-end, 300 base pair reads). The libraries were sequenced to an average depth of ∼22 million unique aligned reads per condition. Two to four technical replicates were used per histone .

### Immunofluorescent staining

MCF-7 cells were seeded on 4-chambered slides (Thermo Fisher, 154917) one day before treatment. DNA-damage positive control cells (corresponding wild-type histone) were treated with 5 mM H_2_O_2_ (Sigma-Aldrich 216763) for 5 min followed by a 30 min recovery in fresh media. The cells were washed three times with PBS, fixed with 4% paraformaldehyde for 15 min at room temperature, and washed three times with PBS. The cells were permeabilized for 10 min using Permeabilization Buffer (PBS containing 0.5% Triton X-100), washed three times with PBS, and blocked for 1 hour at room temperature in PBS containing 1.5% normal goat serum (Thermo Fisher Scientific 50062Z) and 0.1% Tween-20. The cells were incubated overnight at 4°C with phospho-histone H2A.X (Ser139) antibody (Millipore; RRID:AB_309864) diluted at 1:400 in Blocking Solution. The cells were then washed three times with PBS, incubated with Alexa Fluor 488 goat anti-mouse IgG (ThermoFisher; RRID:AB_2534069) diluted at 1:500 in Blocking Solution for 1 hour at room temperature, and washed three more times with PBS. Finally, the coverslips were mounted with Anti-Fade mounting medium with DAPI (Vector Laboratories, H-1200). All images were acquired using an inverted Zeiss LSM 780 confocal microscope (Live Cell Imaging Facility, UT Southwestern Medical Center).

### RNA-sequencing and Data Analysis

#### Generation and sequencing of RNA-seq libraries

MCF-7 cells expressing FLAG-tagged wild-type or mutant histones were seeded in 6-well plates. The cells were collected and total RNA was isolated using the RNeasy kit (Qiagen, 74136) according to the manufacturer’s instructions. The total RNA was then enriched for polyA+ RNA using NEBNext Poly(A) mRNA Magnetic Isolation Module (New England Biolabs, E7490). The polyA+ RNA was then used to generate strand-specific RNA-seq libraries using NEBNext Ultra II Directional RNA Library Prep Kit for Illumina (New England Biolabs, 7760L), barcoded with NEBNext Multiplex Oligos for Illumina (New England Biolabs, E7874L), and sequenced using an Illumina NextSeq 500 to an average depth of ∼40 million reads total per condition. Two independent biological replicates with two sets of sequencing replicates were used.

#### Analysis of RNA-seq data

The quality of RNA-seq datasets was assessed using the FastQC tool (http://www.bioinformatics.babraham.ac.uk/projects/fastqc) and the reads were aligned to the human reference genome (GRCh38/hg38) using Star with default parameters [69]. The uniquely mapped reads were converted into bigWig files for visualization in a UCSC genome browser using writeWiggle function from the groHMM package [70]. Differential gene analysis for WT, H2B and H3 mutants was performed by first counting reads using – featureCounts in the subread package [71] using default parameters. The count matrix was then imported into DESeq2 [72] and differential gene analysis was performed. An FDR cutoff of 0.05 and a fold change cutoff of 1.5 were used to determine significantly regulated genes in mutants compared to the WT H2B and H3 condition.

#### Gene ontology (GO) analyses

Gene ontology analyses were performed on the differentially-expressed gene sets using DAVID (Database for Annotation, Visualization, and Integrated Discovery) [73].

#### Data visualization

Venn diagrams were generated using jvenn [74] for the differentially expressed genes in the different conditions. Heat maps were generated using Java TreeView [75] for genes whose expression was significantly altered in at least one experimental condition.

### Data availability

All raw data and information generated for this study are available upon request from the corresponding authors (W.L.K and D.H.) and the public data repositories noted below. The types of data and information available are as follows:

1. Software, scripts and other information about the analyses can be obtained by contacting the corresponding authors (W.L.K and D.H.).
2. RNA-seq data can be accessed from the NCBI’s GEO database using accession number GSE295718.
3. Genomic sequencing data (raw and processed) for endogenous and ectopic histone genes, as well as the primer pairs used for amplification and sequencing, can be obtained by contacting the corresponding authors (W.L.K and D.H.).

## Author Contributions

A.D.E. – conceptualization, methodology, formal analysis, investigation, writing-original draft, visualization; Y.D. – investigation, validation, visualization; S.S. – investigation, writing-original draft, visualization; M.T. – formal analysis; T.N. – formal analysis; R.K. – methodology, investigation, validation; C.V.C. – methodology, investigation, validation, writing-original draft, writing-review and editing, supervision; D.H. – methodology, investigation, validation, visualization, writing-original draft, writing-review and editing, supervision; W.L.K. – conceptualization, validation, visualization, writing-original draft, writing-review and editing, supervision, project administration, funding acquisition.

## Acknowledgements

We would like to acknowledge Vanessa Schmid at UT Southwestern’s McDermott Center Next Generation Sequencing Core for sequencing the histone gene enrichment in the xenograft tumors, Yoon Jung Kim at UT Southwestern’s Children’s Medical Center Research Institute (CRI) Sequencing Core for the sequencing the RNA-seq sample, and Andrew Lemoff at UT Southwestern’s Proteomics Core for performing the histone mass spectrometry. This work was supported by grants from the NIH/National Institute of Diabetes and Digestive and Kidney Diseases (NIH/NIDDK; R01 DK069710 to W.L.K.), Cancer Prevention and Research Institute of Texas (CPRIT; RP220325 to W.L.K.), and funds from the Cecil H. and Ida Green Center for Reproductive Biology Sciences Endowment to W.L.K.

## Supplementary data

Supplementary data are included with this article. They contain Supplementary Figures S1 through S6.

## Supplementary Figure Legends

**Figure S1. Summary of 236 histone mutations in breast cancer.**

Bar graphs showing the mutation count for each individual mutation (236 total) found in H2A, H2B, H3, and H4. The colored bars represent the top 17 histone mutations selected for further analysis based on mutation count ( ≥2) and amino acid residue of interest (E, R, S, and D). Bar colors indicate H2A (*orange*), H2B (*red*), H3 (*blue*), and H4 (*green*).

**Figure S2. Single nucleotide polymorphisms and pre-existing mutations across breast cancer cell lines.**

Targeted histone gene sequencing was performed on cDNAs generated using a pool of primer pairs that amplifies all 61 core histone genes. Assays were performed in the 8 cell lines indicated.

**(A)** Single nucleotide polymorphisms (SNPs) were identified and the histone gene loci where the silent mutations were found are listed.
**(B)** Missense mutations were identified and listed. The 8 cell lines tested did not harbor the histone mutations of interest, but shared other mutations found in human cancers (*red*). Blue shading indicates that mutations were found in the cell line listed. Darker shading indicates cell lines where the indicated mutation was found at more than one histone gene locus.

**Figure S3. Migration of MCF-7 cells expressing histone mutants.** Cell migration assays were performed in MCF-7 cells ectopically expressing FLAG-tagged histone mutants, as shown in Figure 2A.

(**A-C**) Representative images of migrating MCF-7 cells expressing individual histone H2A (A), H2B (B), and H3 (C) mutants. The cells were stained with crystal violet. Red boxes indicate mutants that had statistically significant effects (refer to Fig. 3B).

**Figure S4. Invasion of MCF-7 cells expressing histone mutants.**

Cell invasion assays were performed in MCF-7 cells ectopically expressing FLAG-tagged histone mutants, as shown in Figure 2A.

(**A-C**) Representative images of invading MCF-7 cells expressing each individual histone H2A (A), H2B (B), and H3 (C) mutant. The cells were stained with crystal violet. Red boxes indicate mutants that had statistically significant effects (refer to Fig. 3D).

**Figure S5. Accumulation of DNA damage in MCF-7 cells expressing histone mutants.** Scatter plots showing quantification of immunofluorescent staining for γH2AX in MCF-7 cells expressing FLAG-tagged histone mutants under basal conditions. Cells expressing cognate wild-type histones treated with 5 mM H_2_O_2_ for 5 min were included as positive controls. The signals in positively stained cells were quantified using ImageJ software and the corrected total cellular fluorescence (CTCF) values were plotted. Each bar represents the mean; n = 3. Asterisks indicate significant differences between mutant and wild-type; One-way ANOVA followed by Dunnett’s multiple comparisons test; * p < 0.05, ** p<0.01, *** p<0.001, **** p<0.0001.

**Figure S6. Summary of phenotypic screens in MCF-7 and MCF-10A cells expressing histone mutants.**

Heatmap summarizing the different phenotypic screens conducted across 17 histone mutants compared to cognate wild-type histones in MCF-7 or MCF-10A cells, and the effects that were observed. Blue indicates phenotypes that were enhanced/increased, green indicates phenotypes that were inhibited/decreased, with the intensity based on significance achieved in each assay compared to its respective wild-type.

