## Supplemental Figures S1-S6 for "Phenotypic and Functional Characterization of Oncohistone Mutations in Breast Cancers"

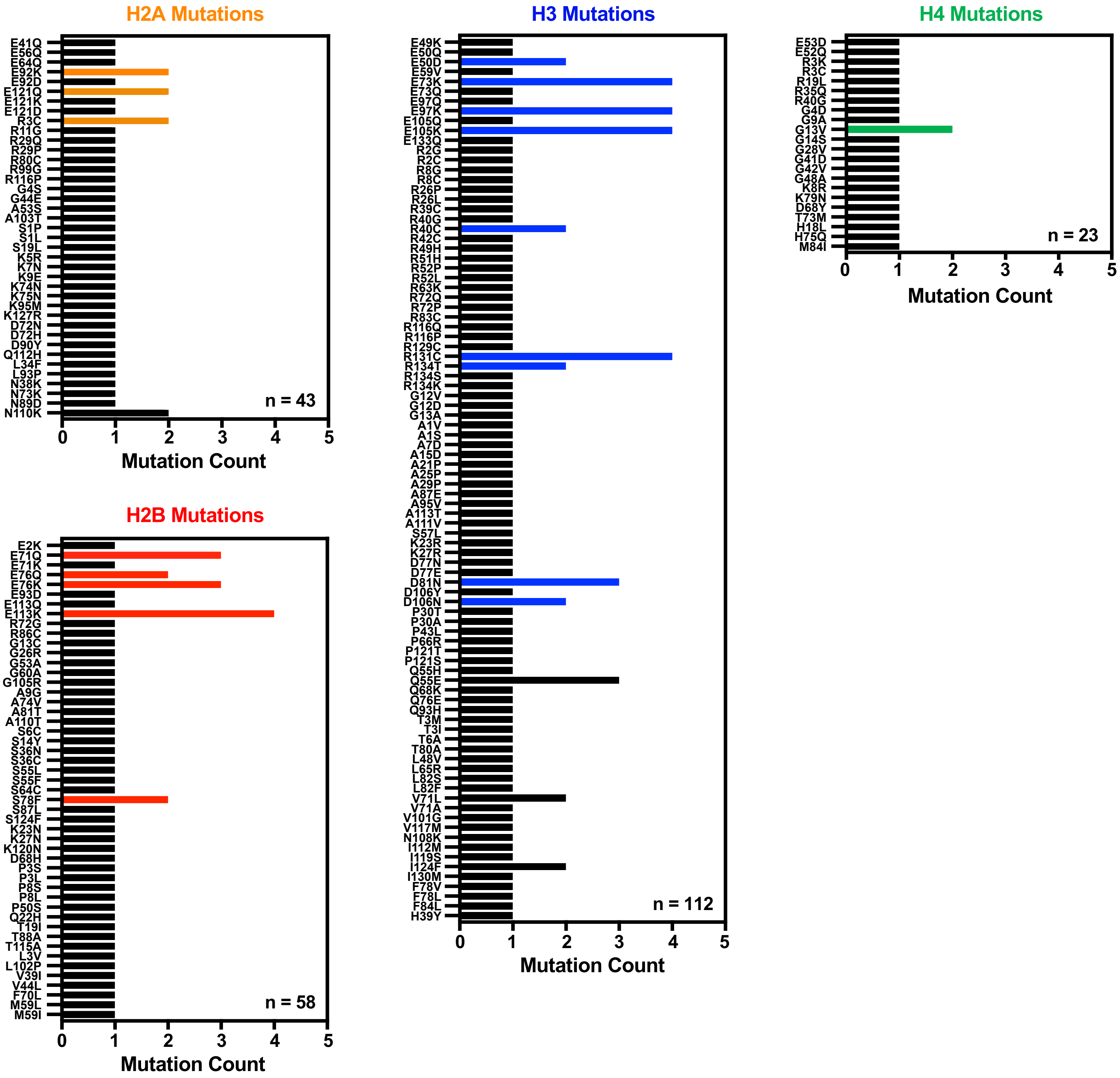
H2A Mutations
E41Q
E56Q
E64Q
E92K
E92D
E121Q
E121K
E121D
R3C
R11G
R29Q
R29P
R80C
R99G
R116P
G4S
G44E
A53S
A103T
S1P
S1L
S19L
K5R
K7N
K9E
K74N
K75N
K95M
K127R
D72N
D72H
D90Y
Q112H
L34F
L93P
N38K
N73K
N89D
N110K
n = 43
0
1
2
3
4
5
Mutation Count
H2B Mutations
E2K
E71Q
E71K
E76Q
E76K
E93D
E113Q
E113K
R72G
R86C
G13C
G26R
G53A
G60A
G105R
A9G
A74V
A81T
A110T
S6C
S14Y
S36N
S36C
S55L
S55F
S64C
S78F
S87L
S124F
K23N
K27N
K120N
D68H
P3S
P3L
P8S
P8L
P50S
Q22H
T19I
T88A
T115A
L3V
L102P
V39I
V44L
F70L
M59L
M59I
n = 58
0
1
2
3
4
5
Mutation Count
H3 Mutations
E49K
E50Q
E50D
E59V
E73K
E73Q
E97Q
E97K
E105Q
E105K
E133Q
R2G
R2C
R8G
R8C
R26P
R26L
R39C
R40G
R40C
R42C
R49H
R51H
R52P
R52L
R63K
R72Q
R72P
R83C
R116Q
R116P
R129C
R131C
R134T
R134S
R134K
G12V
G12D
G13A
A1V
A1S
A7D
A15D
A21P
A25P
A29P
A87E
A95V
A113T
A111V
S57L
K23R
K27R
D77N
D77E
D81N
D106Y
D106N
P30T
P30A
P43L
P66R
P121T
P121S
Q55H
Q55E
Q68K
Q76E
Q93H
T3M
T3I
T6A
T80A
L48V
L65R
L82S
L82F
V71L
V71A
V101G
V117M
N108K
I112M
I119S
I124F
I130M
F78V
F78L
F84L
H39Y
n = 112
0
1
2
3
4
5
Mutation Count
H4 Mutations
E53D
E52Q
R3K
R3C
R19L
R35Q
R40G
G4D
G9A
G13V
G14S
G28V
G41D
G42V
G48A
K8R
K79N
D68Y
T73M
H18L
H75Q
M84I
n = 23
0
1
2
3
4
5
Mutation Count

**A**

| | Human Breast Cancer Cell Lines | | | | | | | |
|---|---|---|---|---|---|---|---|---|
| | Normal | Luminal | | | Triple Negative | | | |
| | Epithelial | Luminal A | Luminal A | HER2+ | Ductal Carcinoma | Claudin Low | Basal | Basal |
| **Silent Mutation** | MCF-10A | ZR-75-1 | MCF-7 | SK-BR-3 | HCC70 | MDA-MB-231 | HCC1937 | MDA-MB-468 |
| H2A R3R;*gene* | | | H2A R3R; *H2AC21* | H2A R3R; *H2AC21* | | H2A R3R; *H2AC21* | | |
| H2A E92E;*gene* | | H2A E92E; *H2AW* | | | | | | |
| H2B E71E;*gene(s)* | H2B E71E; *H2BC12, H2BC11* | H2B E71E; *H2BC12, H2BC11* | H2B E71E; *H2BC12* | H2B E71E; *H2BC12, H2BC17* | H2B E71E; *H2BC12, H2BC8, H2BC18* | H2B E71E; *H2BC12* | H2B E71E; *H2BC12, H2BC17* | H2B E71E; *H2BC12* |
| H3 E73E;*gene* | H3 E73E; *H3C3* | | | | | | | H3 E73E; *H3C3* |
| H3 R134R;*gene* | | | | | | | | H3 R134R; *H3C12* |

**B**

| | | Human Breast Cancer Cell Lines | | | | | | | | |
|---|---|---|---|---|---|---|---|---|---|---|
| | | Normal | Luminal | | | Triple Negative | | | | |
| | | Epithelial | Luminal A | Luminal A | HER2+ | Ductal Carcinoma | Claudin Low | Basal | Basal | |
| | Mutation | MCF-10A | ZR-75-1 | MCF-7 | SK-BR-3 | HCC70 | MDA-MB-231 | HCC1937 | MDA-MB-468 | Percentage |
| H2A | H2A R18C | | | | | | | | | 9 |
| | H2A A41G | | | | | | | | | 9 |
| | H2A L52M | | | | | | | | | 27 |
| | H2A L84F | | | | | | | | | 55 |
| H2B | H2B E3D | | | | | | | | | 36 |
| | H2B D3E | | | | | | | | | 9 |
| | H2B L4P | | | | | | | | | 9 |
| | H2B G27S | | | | | | | | | 55 |
| | H2B I40M | | | | | | | | | 9 |
| | H2B I40V | | | | | | | | | 73 |
| | H2B S76G | | | | | | | | | 82 |
| | H2B G76S | | | | | | | | | 27 |
| | H2B I90T | | | | | | | | | 9 |
| | H2B R93S | | | | | | | | | 9 |
| H3 | H3 R18S | | | | | | | | | 18 |
| | H3 M72I | | | | | | | | | 100 |
| | H3 M72V | | | | | | | | | 100 |
| | H3 V72I | | | | | | | | | 18 |
| | H3 A112V | | | | | | | | | 9 |
| H4 | H4 V3A | | | | | | | | | 18 |
| | H4 R4G | | | | | | | | | 9 |
| | H4 D69H | | | | | | | | | 9 |
| | H4 T72S | | | | | | | | | 9 |

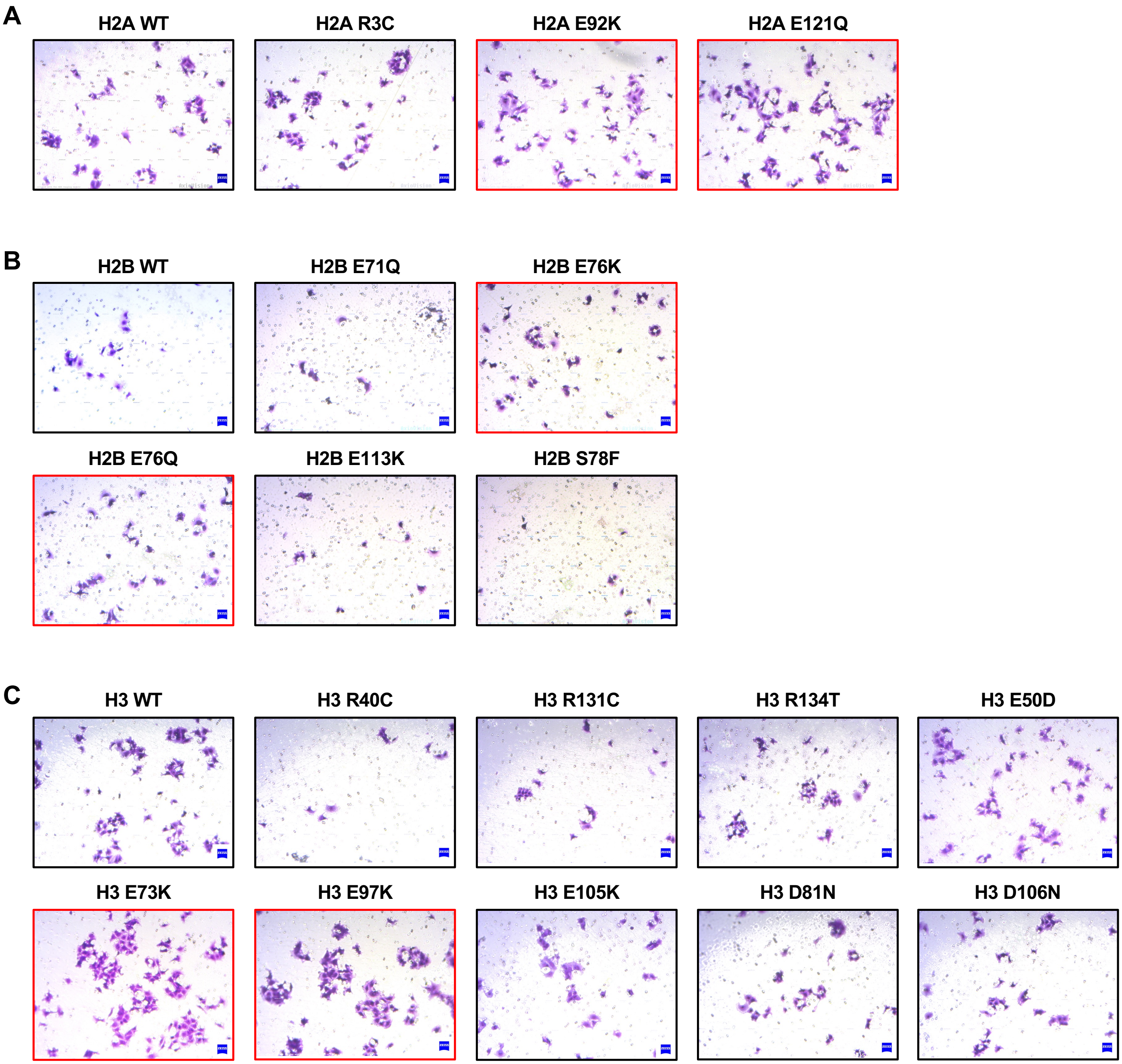
A
H2A WT
H2A R3C
H2A E92K
H2A E121Q
B
H2B WT
H2B E71Q
H2B E76K
H2B E76Q
H2B E113K
H2B S78F
C
H3 WT
H3 R40C
H3 R131C
H3 R134T
H3 E50D
H3 E73K
H3 E97K
H3 E105K
H3 D81N
H3 D106N

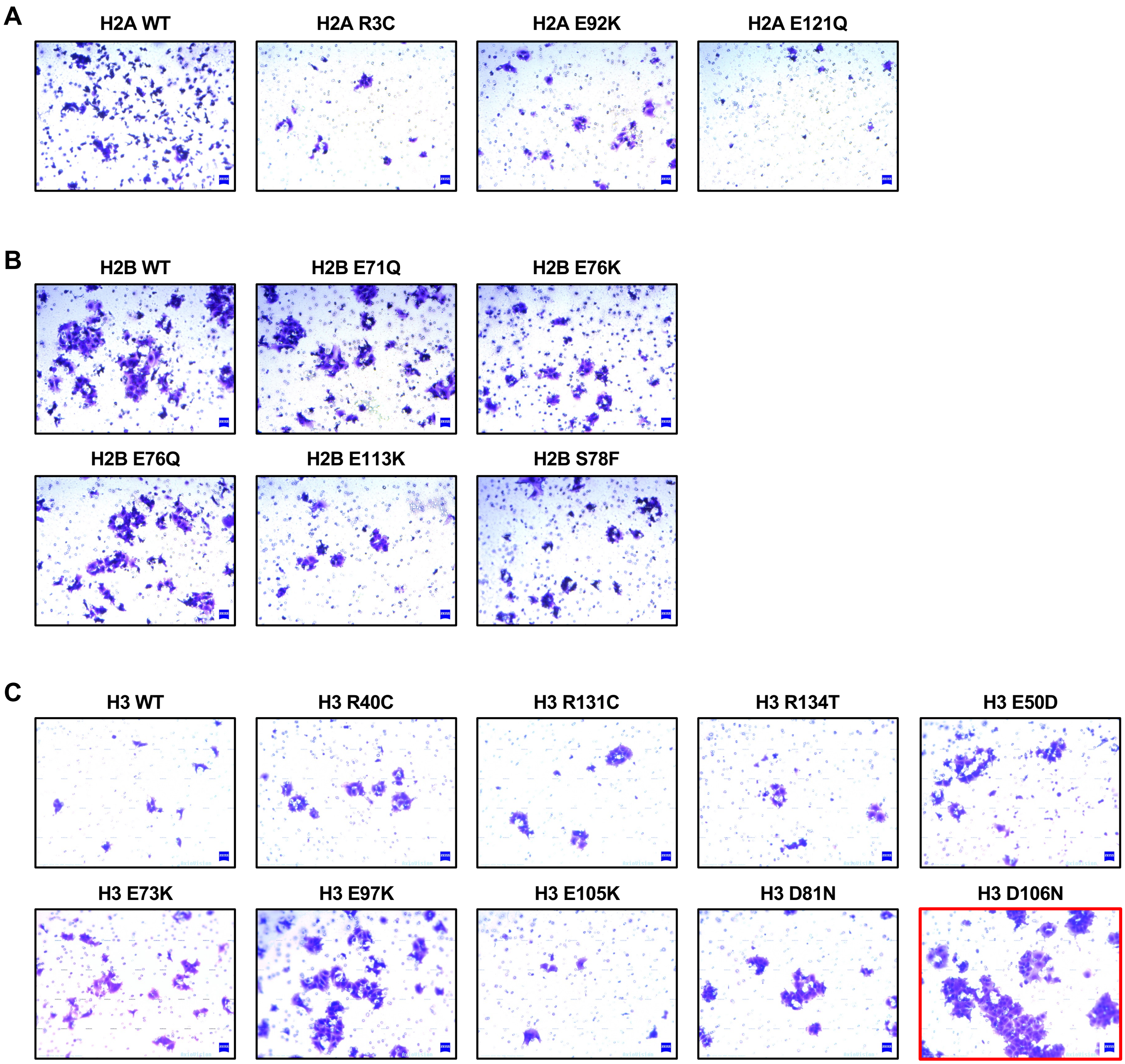
A
H2A WT
H2A R3C
H2A E92K
H2A E121Q
B
H2B WT
H2B E71Q
H2B E76K
H2B E76Q
H2B E113K
H2B S78F
C
H3 WT
H3 R40C
H3 R131C
H3 R134T
H3 E50D
H3 E73K
H3 E97K
H3 E105K
H3 D81N
H3 D106N

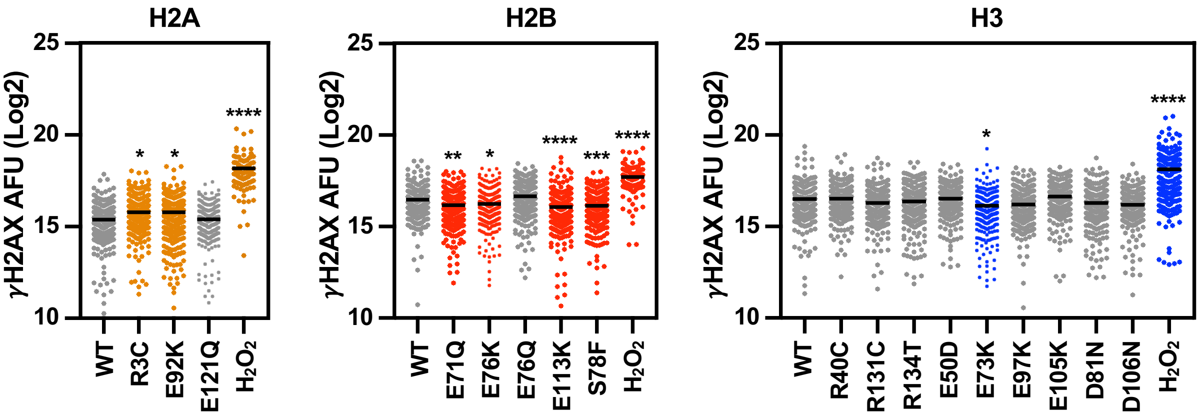
H2A
H2B
H3
γH2AX AFU (Log2)
25
20
15
10
*
*
****
**
*
****
***
****
*
****
WT
R3C
E92K
E121Q
$H_2O_2$
E71Q
E76K
E76Q
E113K
S78F
R40C
R131C
R134T
E50D
E73K
E97K
E105K
D81N
D106N

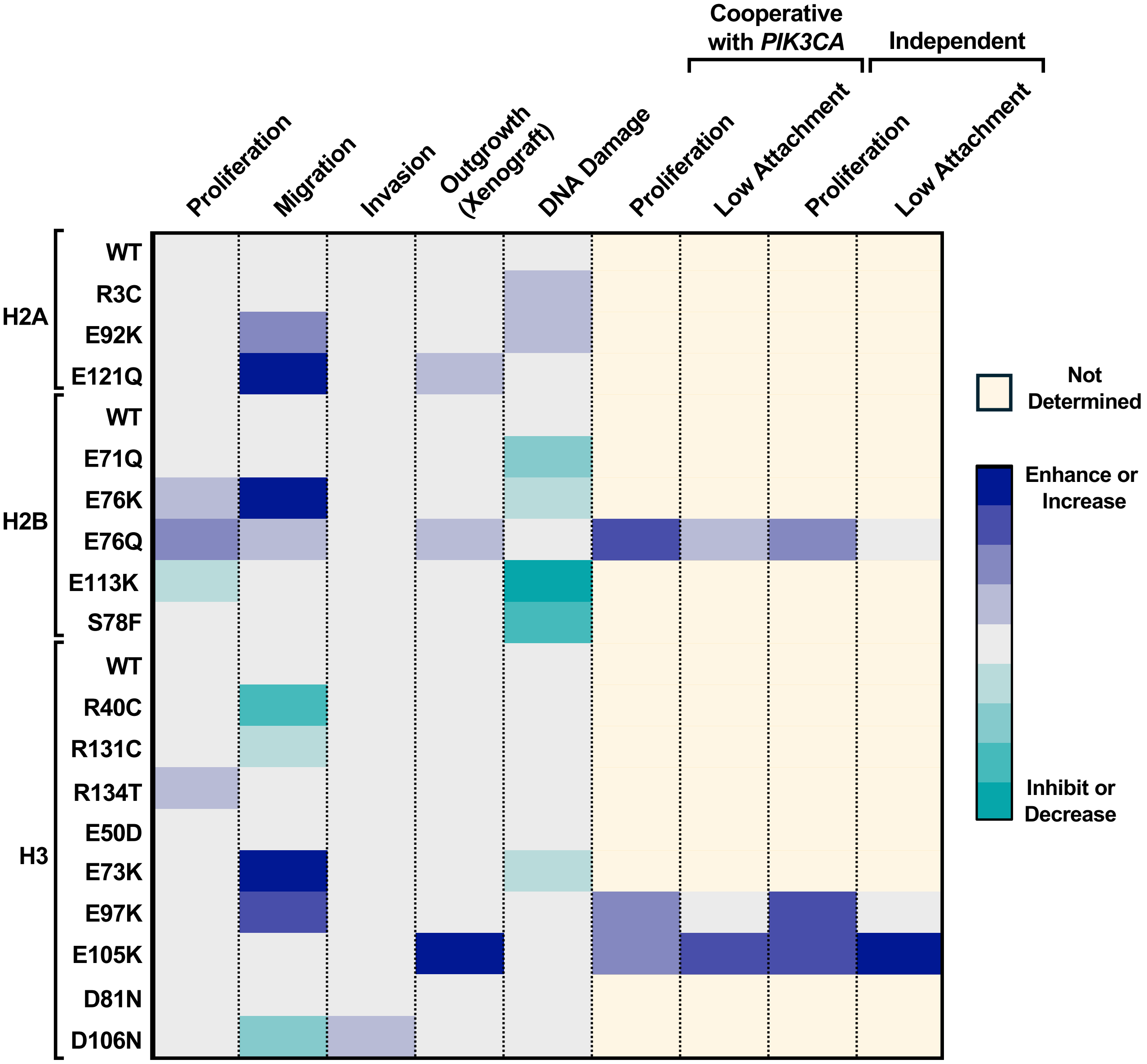
Cooperative with *PIK3CA*
Independent
Proliferation
Migration
Invasion
Outgrowth (Xenograft)
DNA Damage
Proliferation
Low Attachment
Proliferation
Low Attachment
H2A
WT
R3C
E92K
E121Q
H2B
WT
E71Q
E76K
E76Q
E113K
S78F
H3
WT
R40C
R131C
R134T
E50D
E73K
E97K
E105K
D81N
D106N
Not Determined
Enhance or Increase
Inhibit or Decrease
